# Computational comparison of developmental cell lineage trees by alignments

**DOI:** 10.1101/577809

**Authors:** Meng Yuan, Xujiang Yang, Jinghua Lin, Xiaolong Cao, Feng Chen, Xiaoyu Zhang, Zizhang Li, Guifeng Zheng, Xueqin Wang, Xiaoshu Chen, Jian-Rong Yang

## Abstract

The developmental cell lineage tree, which records every cell division event and the terminal developmental state of each single cell, is one of the most important traits of multicellular organisms, as well as key to many significant unresolved questions in biology. Recent technological breakthroughs are paving the way for direct determination of cell lineage trees, yet a general framework for the computational analysis of lineage trees, in particular an algorithm to compare two lineage trees, is still lacking. Based on previous findings that the same developmental program can be invoked by different cells on the lineage tree to produce highly similar subtrees, we designed *D*evelopmental C*e*ll *L*ineage *T*ree *A*lignment (DELTA), an algorithm that exhaustively searches for lineage trees with phenotypic resemblance in lineal organization of terminal cells, meanwhile resolving detailed correspondence between individual cells. Using simulated and nematode lineage trees, we demonstrated DELTA’s capability of revealing similarities of developmental programs by lineal resemblances. Moreover, DELTA successfully identifies gene deletion-triggered homeotic cell fate transformations, reveals functional relationship between mutants by quantifying their lineal similarities, and finds the evolutionary correspondence between cell types defined non-uniformly for different species. DELTA establishes novel foundation for comparative study of lineage trees, much like sequence alignment algorithm for biological sequences, and along with the increase of lineage tree data, will likely bring unique insights for the myriads of important questions surrounding cell lineage trees.

## INTRODUCTION

Life of multicellular organisms typically starts from a zygote, which undergoes multiple rounds of cell divisions and simultaneous differentiation, and eventually develops into an individual organism with multiple types of cells. The developmental *c*ell *l*ineage *t*ree (CLT) is a record of both the differentiation states or types of every single cell at a specific developmental time point, and all the cell division events since the zygote that lead to these cells (Fig. 1A). More generalized CLT does not necessarily root at the zygote, but may start from any dividable cell, which is a subtree of the CLT rooted at the zygote, or sub-CLT (Fig. 1A). As one of the most important traits of multicellular organisms, CLT is the key to many significant unresolved problems in life sciences. For example, developmental CLT records the details of developmental process^1–6^, and helps to explain the mechanism of developmental robustness at the cellular level^7–9^. Other types of CLT reveal the origin of relapsed or metastatic tumor cell population ^10,11^, the risk of carcinogenesis attributable to the number of cell divisions since the zygote ^12,13^, as well as the origin and evolution of cell types and lineages ^14,15^.

**Fig. 1.**
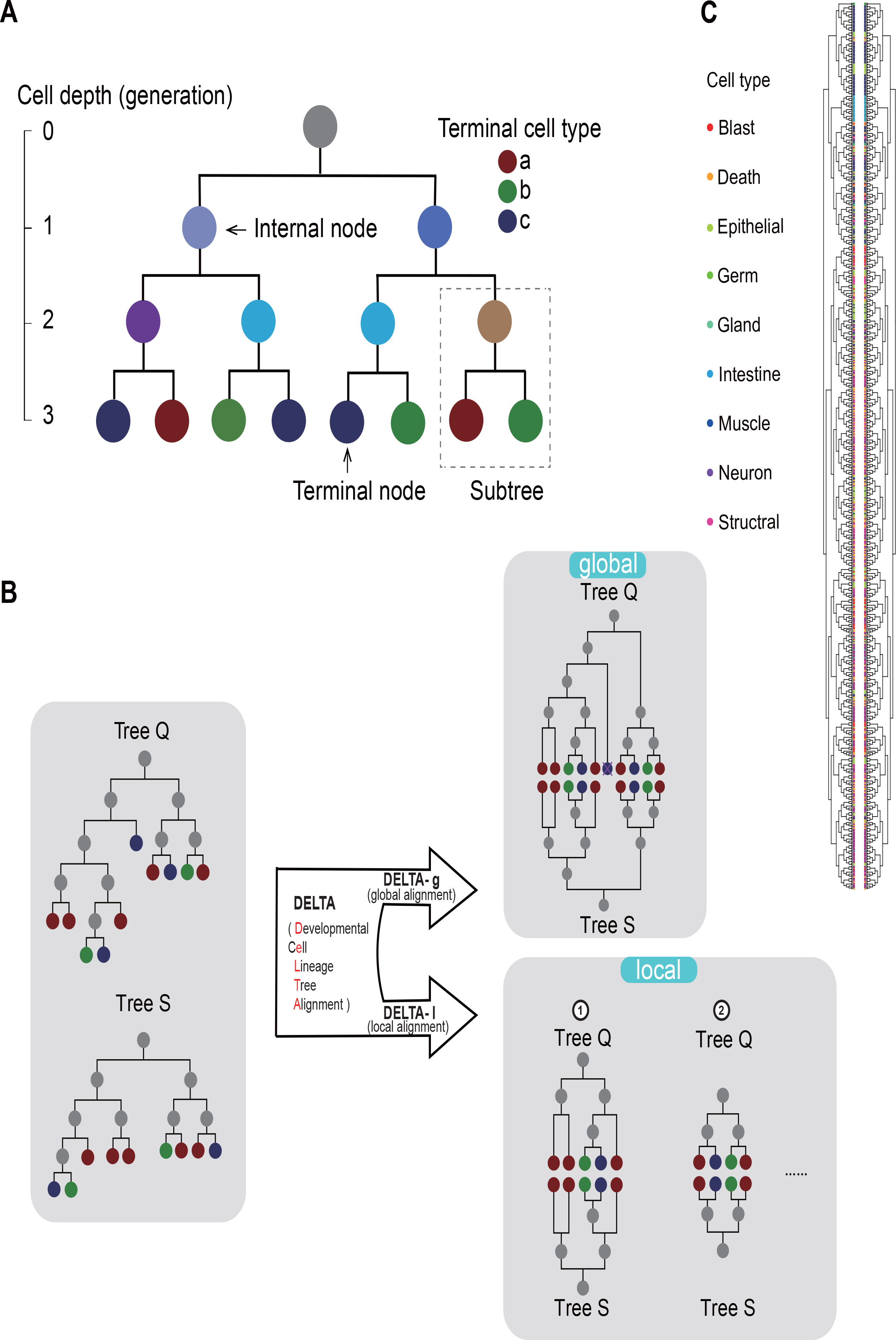
Overview of DELTA algorithm. (A) A very simple typical developmental cell lineage tree (CLT) rooted at the zygote (“Z”). Cells undergo further division were represented by internal nodes, and others terminal nodes. Cell types of the terminal nodes were indicated by the color legend on the right, whereas the (epi-)genetic state of an internal cell, as inferred from the cell type of the two daughter cells, were also indicated by different node colors. The depth, or the number of divisions since the zygote, of a cell is indicated by the vertical axis. A subtree, or sub-CLT, is outlined by a dotted box. (B) Two CLTs, *Q* and *S*, with types of their terminal cells were color-labeled. DELTA aligns them globally (DELTA-g), where all cells in respective CLTs are either pruned or aligned, or locally (DELTA-l), where only pairs of sub-CLTs with good enough alignments were reported (See Fig. S1A for more details). (C) The DELTA alignment of the *Caenorhabditis elegans* CLT of standard anatomical terminal cell type annotation, with an isomorphic version of itself, where 30% randomly chosen sister sub-CLT pairs were swapped. The resulting CLT alignments were visualized by our newly developed R package, “ggVITA” (See also Fig. S1C).

More than 35 years ago, the seminal work of John Sulston and colleagues in the nematode *Caenorhabditis elegans* had revealed the first complete developmental CLT containing ~1000 terminal cells via painstaking observation through optical microscope ^16^. Advancement of microscopic imaging techniques had since facilitated assessments of developmental CLT in other nematodes ^17–20^ and ascidian ^21^, but is of limited help in more complex species where the body is opaque and consists of vastly more cells. Recently, the application of genome editing in combination of single-cell high-throughput sequencing ^1,2,4,22,23^ has open a new path towards the CLT in more complex species, where the topology of the CLT is determined by phylogenetic analysis informed by the genomic editions, and the cell types by transcriptome profiles of single cells. Although it has yet to achieve the resolution of individual division events and types for all terminal cells, this new technique will likely further improve and generate more and better CLT data in the near future due to the scientific significance of CLT.

Unfortunately, a general computational framework for alignment between CLT is currently lacking. Here alignment means finding fine-scale similarities between two structured entities. As an analogy, comparison between two biological sequences, such as DNA or protein, can be accomplished by alignment. Indeed, the development of sequence alignment algorithm in the 1970s ^24,25^, has built the foundation for the interpretation of biological sequences, way before the soar of sequencing data. Similarly for CLTs, there are also common research needs that might be fulfilled by a proper alignment algorithm. First, comparison of relevant data is required for quality assessment. For example, sequence/CLT alignment can evaluate how replicable the results from two technical repeats are. Second, comparison is required to relate new observations to the known data, increasing the interpretability of new observations, such as a newly determined sequence subjected to homology search for an initial guess of its function or source. Third, comparison between individual observations is required to disentangle variations from the consensus, and to ultimately find the underlying law constraining the consensus and generating variation. Last but not least, comparative analysis, which by definition is comparison between sequences/CLTs from different species, is a major way of studying evolution, the ultimate causation of many biological patterns. Currently, these obstacles are only partially addressed by comparing relative fraction of different types of cells generated by CLTs. However, just as in sequence comparison, majority of information is loss when fraction of A/T/G/C is compared, relative to when sequence is aligned, finding the fine-scale alignment between CLTs will likely reveal much more biological information.

To address this critical demand, we designed DELTA (*D*evelopmental C*e*ll *L*ineage *T*ree *A*lignment), an algorithm that aligns a pair of CLTs by identifying homeomorphic sub-CLTs, with the assumption that similar genetic (or developmental) programs should give rise to similar sub-CLTs ^7,26^. Using simulated CLTs ^27^ and the real CLTs from *C. elegans* ^28^, we showed that homeomorphic sub-CLTs found by DELTA have highly similar expression profiles. Comparison among CLTs of wildtype and single gene knock-out mutants of *C. elegans* ^3^ revealed both known ^5^ and novel homeotic transformations of cell fates in the mutant strains, and suggested for the deleted genes functional relationship compatible with evolutionary and experimental evidence. Finally, we compared developmental CLTs of two nematodes and pinpointed evolutionary changes of fates between cells on these two CLTs. By maximizing the alignment score between real CLTs of the two species, we found biologically interpretable correspondence between their non-uniformly defined cell types, highlighting a conceptually new way of finding the evolutionary relationship between cell types. Just as sequence alignment algorithm had fundamentally transformed genetics, the possibility of CLT comparison/alignment opened by DELTA will likely lead to new opportunities for deeper understandings of biology of multicellular organisms, such as assessing the repeatability of differentiation, linking sub-CLTs to developmental programs, and distinguishing autonomous and regulatory components in development.

## RESULTS

### Overview of DELTA algorithm

A typical developmental CLT as analyzed here is a binary tree (Fig. 1A), where each node represents a single cell and each branch represents a descendant relationship pointing from a mother cell to one of its daughter cells. The cells in the tree can be divided into internal or terminal cells/nodes by whether they undergo further division as recorded by the CLT. A subtree rooted at any of the cells is a sub-CLT. The terminal cells of the CLT are all labeled by their cell types, which could be anatomically defined as, for example, muscle, neuron, or defined by the expression state of one or more genes such as CD4+ cells. Note that unlike CLTs commonly discussed in nematodes such as *C. elegans*, we ignored the temporal duration of cell cycle and lineal order of sister cells to ensure compatibility to CLTs determined by genomic barcoding. In other words, the length of branch hold no information about how long each single cell exists, and swapping any pair of sister sub-CLTs (i.e. two sub-CLTs whose roots are a pair of sister cells divided from the same mother) will not change the CLT.

We designed the *D*evelopmental C*e*ll *L*ineage *T*ree *A*lignment (DELTA) algorithm with the purpose of identifying similarities in developmental programs using the phenotypic information represented by the CLT. Here the developmental program means a succession of cell fate choices made at every division events recorded by the CLT. We assumed the developmental state of a cell is reflected by those of its daughter cells, which is further defined by their daughter cells thereinafter until the terminal cells with known cell types (Fig. 1A, color of nodes). In other words, a pair of cells is similar if the two sub-CLTs rooted at them resemble each other in topology and lineal organization of terminal cell types. This assumption was deemed useful in demonstrating the simplicity ^26^ and robustness ^7^ of metazoan CLTs, as well as identifying homeotic transformation of cell fates ^5^. DELTA compares every sub-CLTs from a query CLT with those from a subject CLT, and exhaustively search for their maximal resemblance in topology and lineal organization of terminal cell types via a dynamic programming strategy (Fig. S1A). As an analogy, sequence alignment algorithms align residues in biological sequences with the constraint of their sequential order, whereas DELTA aligns terminal and internal cells in CLTs with the constraint of their lineal organization. DELTA can align CLTs globally (DELTA-g), where all cells in respective CLTs are either pruned or aligned, or locally (DELTA-l), where only pairs of sub-CLTs with good enough alignments were reported, much like global and local sequence alignment, respectively (Fig.1B). DELTA also estimates statistical significance of each CLT alignment relative to random pairs of CLTs with same sizes and terminal cell type compositions as the aligned CLTs. More algorithmic details of DELTA are given in Methods and Supplementary Text S1.

To validate the correctness of our DELTA implementation, we aligned *C. elegans* CLT with an isomorphic version of itself, where 30% randomly chosen sister sub-CLT pairs were swapped. DELTA-g successfully aligned the isomorphic CLT with the original by matching all terminal nodes, yielding a DELTA score same to that of the alignment between two identical *C. elegans* CLTs (Fig. 1C. See also Fig. S1B). We also developed an accompanying R package named ggVITA (*gg*tree-based *vi*sualization of *t*ree *a*lignments) for the visualization of DELTA alignments (Fig. 1C. See also Fig. S1C).

### CLT simulations suggest DELTA can identify developmental similarity

To further demonstrate that DELTA alignment can indeed reveal developmental similarities, we simulated CLTs using a previously published model ^27^, in which the epigenetic on/off state of each gene in each cell and each discrete time point was calculated by a predefined regulatory network. (Fig. 2A and B. See Methods) DELTA-l was used to align the simulated CLT with itself. This process was repeated with 1,000 different simulated CLTs, and the top ten local alignments from each simulation were examined to assess the performance of DELTA. Several results in support of the capability of DELTA were observed. First, we found that for each simulated CLT, the self-alignment always has the highest DELTA score in the local alignment result (Fig. 2C, left most red dot). Second, DELTA tends to find alignments between large sub-CLTs, which contain more developmental information (Fig. 2D, red dots). Third, the CLT alignments are statistically highly significant, indicating DELTA scores are much higher than those between random CLTs of similar sizes and terminal cell type compositions (Fig. 2E, red dots). Fourth, the epigenetic distance (measured by hamming distance) between the roots of the aligned sub-CLTs is much smaller than that of two randomly chosen internal cells, and tends to be lower for those with higher DELTA scores (Fig. 2F, red dots). Fifth, by comparing the expression (i.e. epigenetic on/off state) of each gene for all aligned (terminal and internal) cells in a pair of (sub-)CLTs, we found that their expression trajectories were much more similar than expected (hamming distance normalized to [0,1], with expectation of 0.5) between aligned internal or terminal cells (Fig. 2G), suggesting that not only the initial state, but also the subsequence changes of expression is highly similar between the aligned CLTs. These results demonstrated that a DELTA can indeed pair up internal cells with similar epigenetic states.

**Fig. 2.**
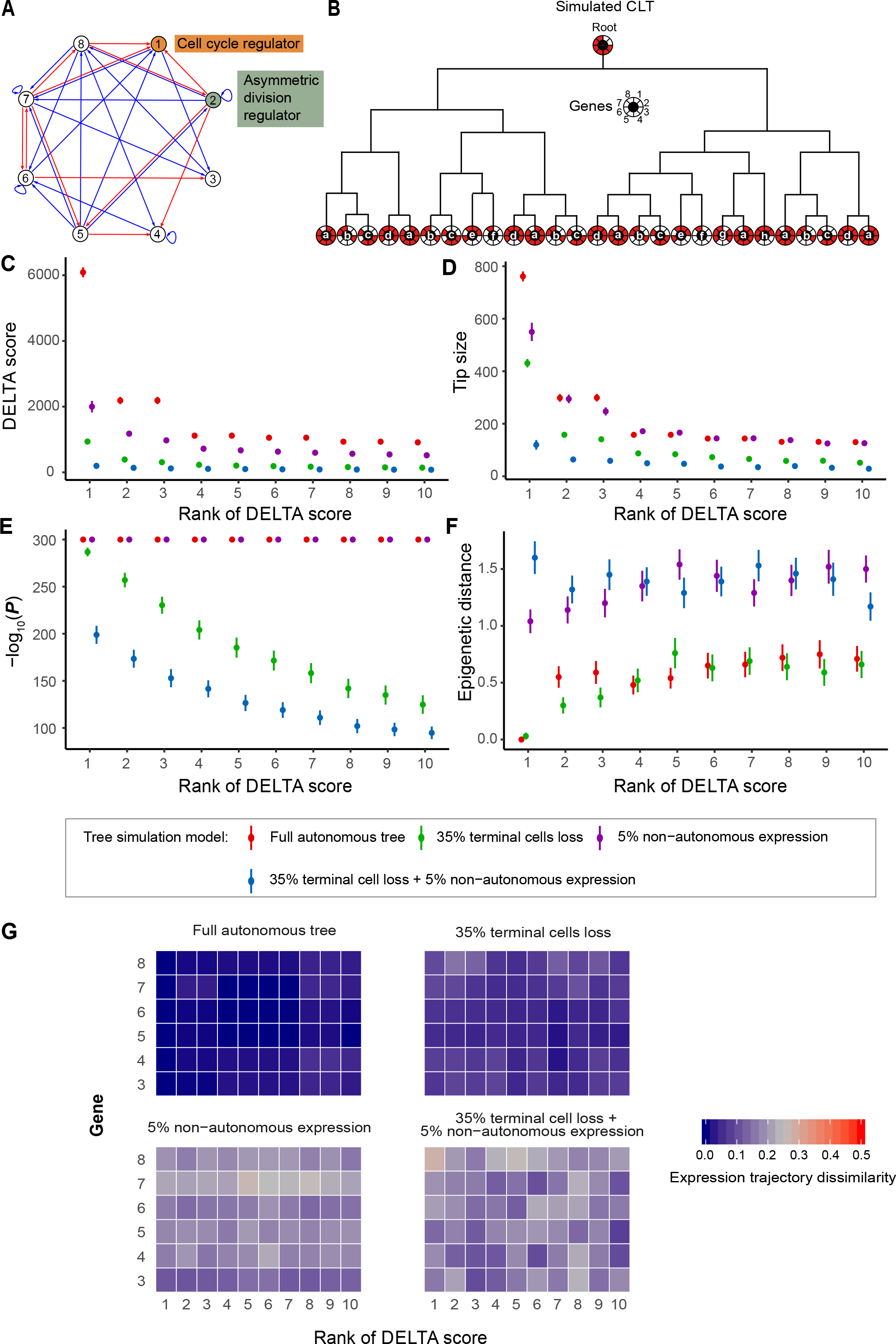
Validating DELTA by simulated CLTs. (A) An example of transcriptional regulatory network used to simulate CLT. There are eight genes each regulated by an average of four other genes. Red and blue lines represent activations and repressions, respectively. Values in the matrix of regulatory interactions are detailed Fig. S2A. (B) An example of simulated cell lineage tree. Lowercase letters a to h represent eight terminal cell types based on the ON/OFF state of eight genes, which is shown as red (ON) or white (OFF) on the corresponding sector around the letter. The development of the lineage tree stops when one of the terminal cells reaches 12 rounds of divisions or the 50^th^ discrete time point of the simulation. CLT were also simulated under other different parameter settings, as listed in Fig. S2B and explained in Methods. (C) The score of the top 10 CLT alignments found by DELTA-l from self-comparison of simulated CLTs. Besides the full simulated CLT (red dots), perturbations were added to mimic experimental/biological noises, such as loss of terminal cells (green dots), non-autonomous cell fate (purple dots), or both (blue dots). Each dot shows the average score and its standard error assessed by 1,000 simulations with different regulatory network and initial expression state. (D-F) Similar to (C), except that the number of terminal cells of the aligned CLT (D), statistical significance of the alignment (E) and epigenetic distance between roots of aligned CLTs as measured by hamming distance of expression states (F) are plotted. (G) For the same set of top 10 CLT alignments presented in (C) (x axis), the expression trajectory dissimilarity of every gene except gene 1 and 2 (y axis) were shown. The expression states of a specific gene at the last time point of every cell were compared for every pair of matched cells from the two aligned CLTs. The resulting hamming distance is normalized by the number of matched cell pairs, giving rise to the expression trajectory dissimilarity, whose value was scaled as the color scale bar to the right. All expression trajectory dissimilarities were averaged values from DELTA-l results of 1,000 simulations. The expression trajectory dissimilarities between the aligned cells are clearly much lower than null expectation of normalized hamming distance = 0.5.

Two practical considerations prompted us to further scrutinize the performance of DELTA using revised models of CLT simulation. First, the aforementioned model used in simulated CLTs implicitly assumed that all cells differentiate autonomously, whereas the real differentiation process is believed as highly regulatory, i.e. affected by external signals from other cells or the environment. We thus introduced a probability (5%) of randomly flipping the epigenetic state of a gene in every time point during the simulated development, by negating its expression level (See Methods), excluding the cell cycle and asymmetric division regulator. Second, current experimental techniques are not perfect in capturing all single cells, making the experimentally reconstructed CLTs incomplete. For example, the state-of-the-art lineage tree reconstruction method ^23^ isolates single cells using the 10x Chromium, an instrument with an average of 35% cell losses. To reflect such technical limitation, we simulated two CLTs with identical initial parameters (regulatory network and expression of the root), but different random loss of 35% terminal cells, reconstructed the simulated CLTs following their actual lineal relationship (See Methods), and aligned them by DELTA. As expected, these two perturbations reduced the DELTA score (Fig. 2C), CLT size (Fig. 2D), statistical significance (Fig. 2E), and the epigenetic similarity between the aligned sub-CLTs (Fig. 2F and G). Nevertheless, it is still capable of identifying statistically significant and epigenetically similar sub-CLTs (Fig. 2C-G). We found by additional simulations that the performance of DELTA remains satisfactory with 5, 10, 20 or 50% cell losses, but not 90% (data not shown). Nevertheless, these result suggested that, despite of the detection power reduction due to the stochastic perturbations, associating CLT phenotypes to underlying epigenetic states by DELTA remains feasible.

### Alignment of *C. elegans* CLT by DELTA reveals cells with highly similar developmental programs

Next, we sought to test the performance of DELTA using real CLTs from *C. elegans*. The *C. elegans* embryonic CLT contains 8 pairs of bilaterally symmetrical sub-CLTs ^16^ with 10 or more terminal cells (Fig. 3A). We aligned these symmetric pairs of sub-CLTs by DELTA-g, and found that their DELTA scores are highly significant (Fig. 3B, gray bars), and are always higher than those from the alignments between one sub-CLT from the symmetric pairs, and another sub-CLT with a similar number of terminal cells (Fig. 3B, blue dots. See Methods). These results showed that DELTA efficiently aligns symmetric pairs of sub-CLTs, which likely have identical developmental states.

**Fig. 3.**
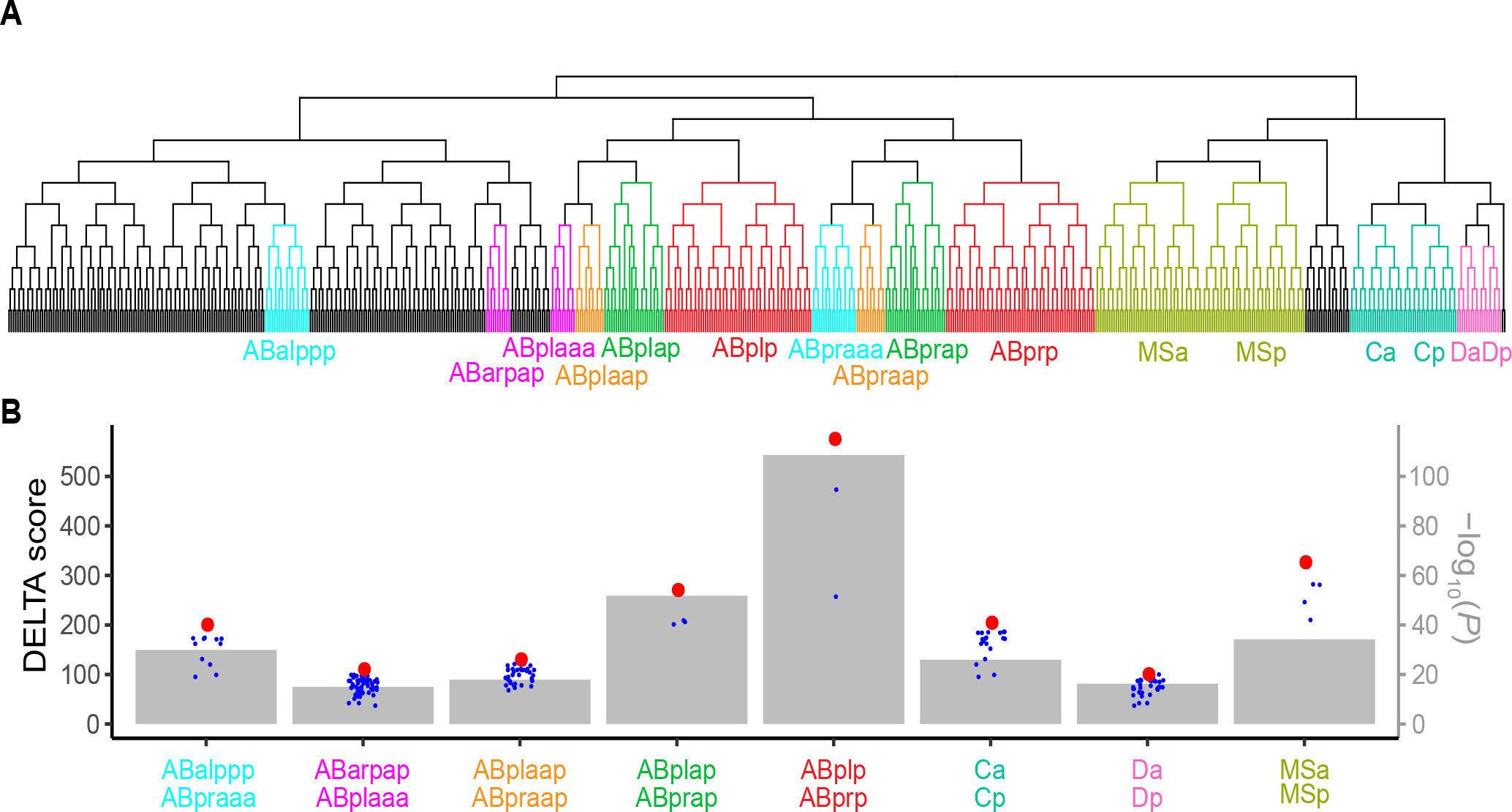
Bilaterally symmetric sub-CLTs yielded highly significant DELTA alignment (A) Bilaterally symmetric sub-CLTs in *C. elegans* CLT were highlighted by different colors, whereas the lineal names of their root were also marked below every sub-CLTs. (B) The alignment score (dots, scaled by the left y axis) and statistical significance (gray boxes, scaled by the right y axis) found by DELTA-g alignment between the symmetric sub-CLTs were indicated by red dots. As controls, sub-CLTs whose number of terminal cells differs from the symmetric sub-CLTs by no more than 10% were also compared to one of the symmetric sub-CLTs, and the resulting DELTA scores were indicated by the blue dots.

To further assess the capability of DELTA in finding similarities in developmental programs, we take advantage of the EPIC database (Expression Patterns in *C. elegans*), where expression of 130 genes were tracked in each cell during the embryonic development of *C. elegans* from the zygote to the larva ^29^. We collected the top 1,000 alignments in DELTA-l results of *C. elegans* CLT *vs* itself, and calculated the Pearson’s correlation coefficient *R* between the aligned cells in the sub-CLTs. Since the sizes of different alignments vary, the Pearson’s *R*s are standardized by Fisher’s r-to-z transformation before being compared. In support of the usefulness of DELTA, higher *z* is observed in CLT alignments with higher DELTA score (Fig. 4A, Pearson’s *R* = 0.951, P<10^−300^, Spearman’s ρ = 0.926, *P* < 10^−300^) and more significant alignment *P* values (Fig. 4B, Pearson’s *R* = 0.504, P < 10^−98^, Spearman’s ρ = 0.176, *P* < 10^−11^). In combination with the results from the simulated CLTs, we demonstrated that DELTA can indeed identify CLTs with highly similar developmental programs.

**Fig. 4.**
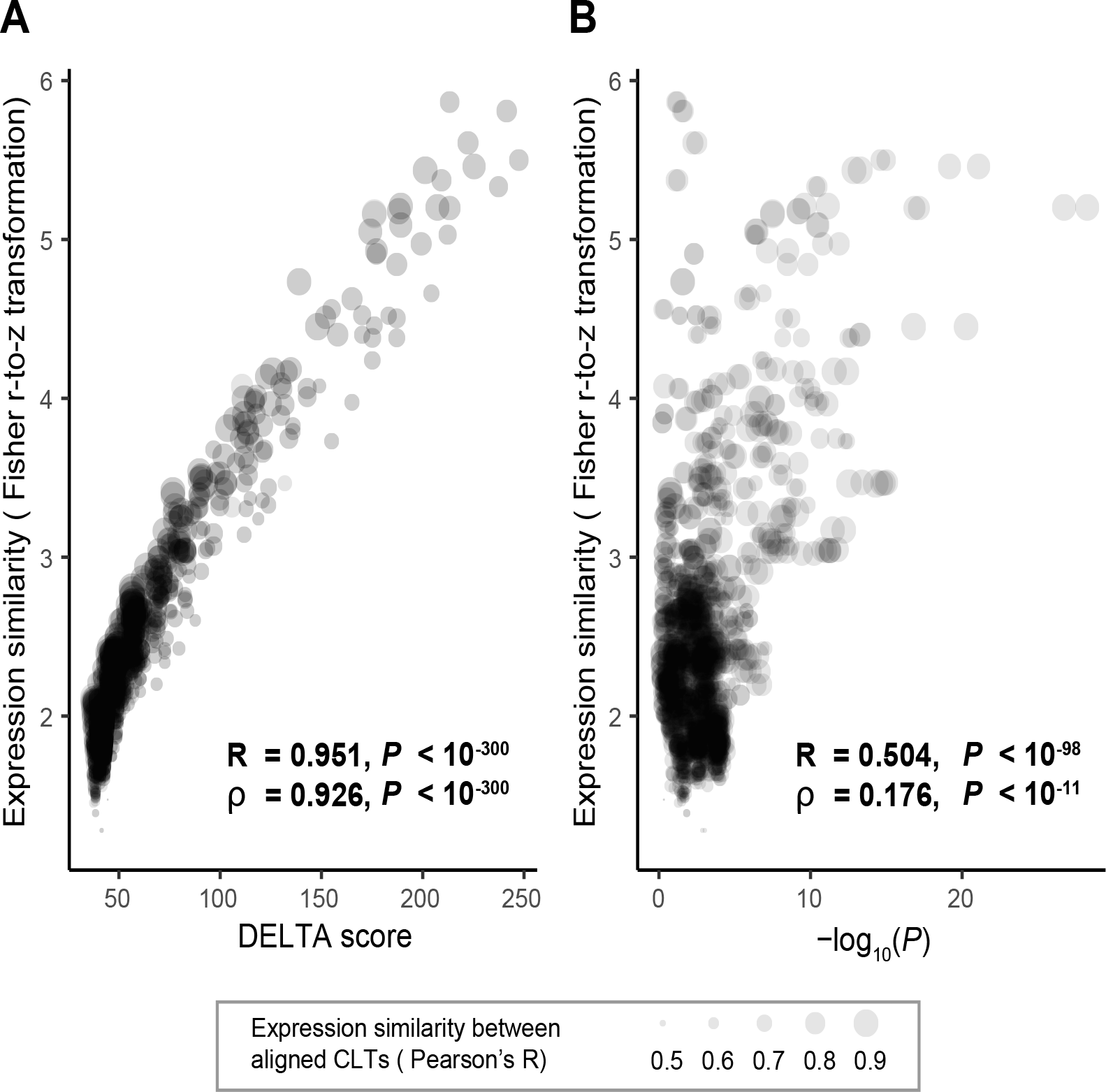
Expression similarities between aligned sub-CLTs in *C. elegans* Top 1,000 local alignments between sub-CLTs of *C. elegans* were found by DELTA-l, and those with nominal *P* < 0.05 were checked for expression similarities between aligned cells. For each sub-CLT alignment, the expression level of 130 genes recorded by EPIC were compared for all aligned cells, giving rise to the expression similarity between the two aligned sub-CLTs in the form of a Pearson’s Correlation Coefficient (point size). The expression similarities were further processed by Fisher’s *r*-to-*z* transformation to ensure comparability between sub-CLT alignments with different number of cells, and then shown to be highly correlated with the DELTA score (A) and *P* value (B) of the sub-CLT alignment.

### Phenotypic changes of mutant CLT quantified by DELTA reveals functional relationship among underlying genes

Inspired by the capability of DELTA in identifying similarities in developmental programs by homeomorphic (sub-)CLTs, we continued to test whether DELTA can associate CLT changes to their underlying genetic mechanisms. The database of Digital Development ^3^, where CLTs are recorded for *C. elegans* strains with ~ 200 conserved genes individually knocked out (KO strains), provides a unique opportunity of comparing phenotypic changes in CLT with underlying genetic differences. Specifically, homeotic transformations, where a cell *x* adopts the fate used by another cell *y* in normal development (an *x*-to-*y* transformation), were previously observed using this dataset ^5^. Here the fate of a cell was indicated by the lineal name of the cell. For a homeotic transformation of *x*-to-*y* in a mutant strain, we extracted the sub-CLT rooted at *x* from the mutant strain, as well as the sub-CLT rooted at *y* from the wildtype strain. Using a scoring matrix defined by the number of markers with shared expression (Fig. 5A) and a prune cost of 1, we used DELTA-g to align all the extracted pairs of sub-CLTs from the homeotic transformations, i.e. the sub-CLT with an altered fate in the mutant strain, and the sub-CLT from the wildtype strain representing the transformed new fate. We found that 93.5% of them gave rise to statistically significant (*P* < 0.05) alignments (Fig. 5B and C), suggesting that DELTA can indeed identify homeotic cell fate transformations. Moreover, DELTA elaborated the correspondence between the terminal cells of these aligned CLTs (Fig. 5D, alignments on the left), revealing the subtle differences between wildtype and transformed sub-CLTs.

**Fig. 5.**
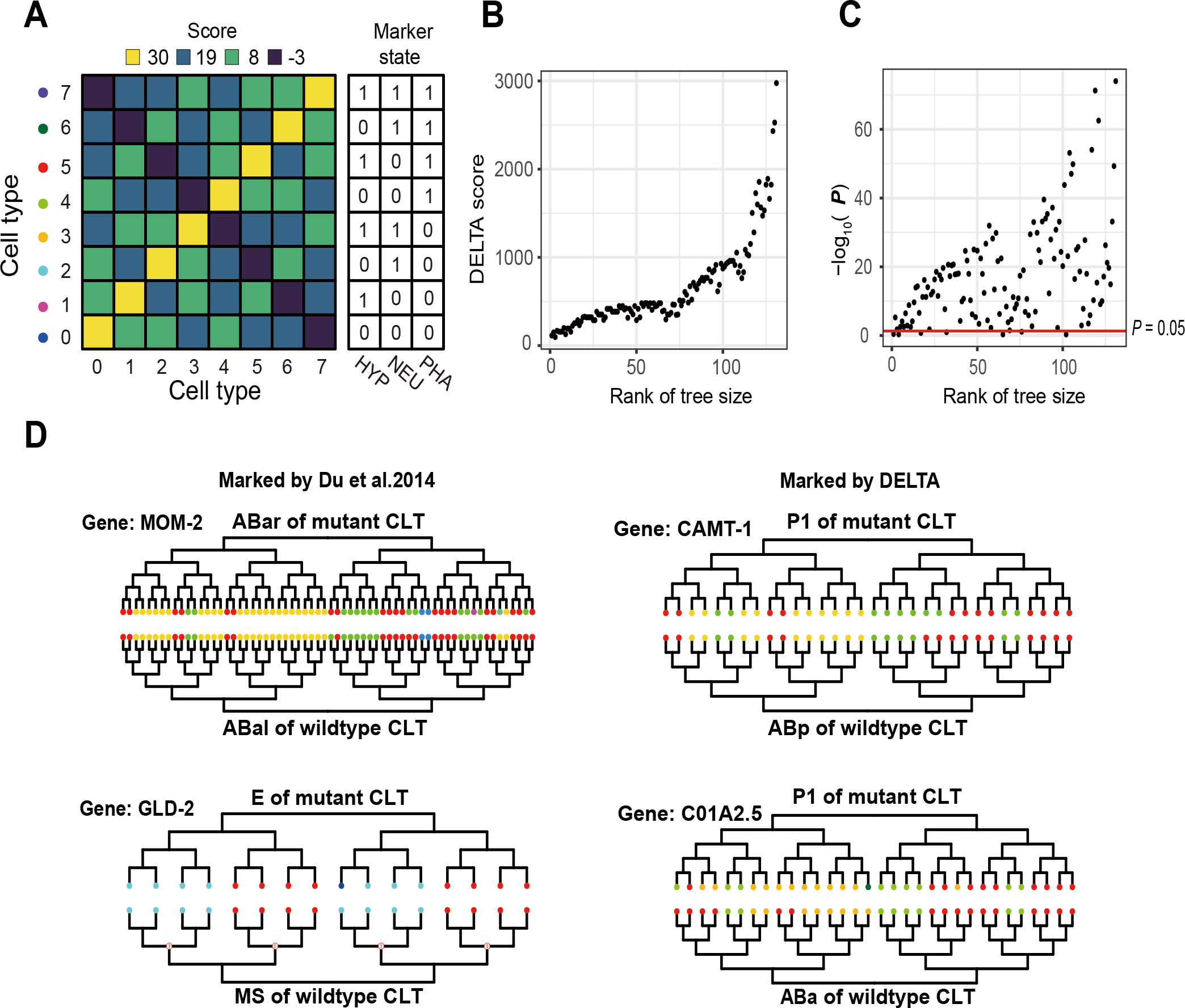
DELTA reveals homeotic cell fate transformation in *C. elegans* mutant (A) The scoring matrix used in DELTA analyses for cell types annotated in the Digital Development database. (B-C) For the 131 homeotic cell fate transformation events found by Du et al. between CLTs with at least 5 terminal cells, the DELTA score (B) and *P* value (C) between the sub-CLTs in mutant strains and that in wildtype strain representing the adopted cell fate were shown. (D) Detailed sub-CLT alignments visualized by our newly developed R package “ggvita”. The two alignments on left were previously marked by Du et al., and the two on right were newly found by DELTA. Terminal cell types are indicated by colors same as in (A).

We further looked the top 100 DELTA-l results between wildtype and each mutant strain for homeotic transformations events. Some known homeotic transformations are among the top ranking local alignments. For example, the cell fate transformation of ABar lineage in MOM-2 KO strain into ABal in wildtype strain has the 6th highest DELTA score in local alignment between wildtype and MOM-2 KO CLT, whose detailed alignment between individual cells found by DELTA were visualized by ggVITA (Fig. 5D, top left alignment). Similarly, the E cell in GLD-2 KO strains takes the cell fate of wildtype MS, which corresponds to 32^nd^ top alignments in DELTA-l results between wildtype and GLD-2 KO CLTs (Fig. 5D, bottom left alignment). Furthermore, we found some alignments between sub-CLTs that likely correspond to addition homeotic cell fate transformation that is not previously reported, such as the transformation of P1 into ABp when CAMT-1 is deleted, and P1 into ABa when C01A2.5 is deleted (Fig. 5D, right alignments). Note, however, that most local alignments found in DELTA-l between wildtype *vs* mutant strains are between sub-CLTs that is unchanged by the gene knockout, or sub-CLTs that are highly similar in the original CLT. Nevertheless, these results suggest that homeotic cell fate transformation can be readily found by DELTA.

Given the above result, we further hypothesized that the phenotypic impact of the gene deletions, as approximated by the DELTA score between wildtype and KO CLTs, can reflect the functional importance of the underlying genes. To test this hypothesis, we compared the DELTA score from global alignment between wildtype and KO CLTs with the evolutionary rate of the genes being knocked out, since functionally more important genes generally evolve more slowly and thus are more conserved ^30^. Here, we used the dN/dS ratio to measure protein evolutionary rate, where dN is the number of nonsynonymous nucleotide substitutions per nonsynonymous site, and dS is the number of synonymous nucleotide substitutions per synonymous site ^31^. We split the genes with KO CLTs into two groups with high or low DELTA score with the wildtype CLT and compared the average evolutionary rate of the two groups. As we divided the two groups by greater DELTA score differences, the deviation of evolutionary rate of the two groups continues to increase to up to ~ 8 fold when genes with DELTA score > 7400 and < 2600 were compared (Fig. 6A). Since genes with more dramatic functional impact upon deletion are generally more constrained by natural selection ^30^, this observation suggests that DELTA comparison between KO and wildtype CLT can indeed quantify phenotypic changes in CLT in relation to the functional importance of the deleted gene.

**Fig. 6.**
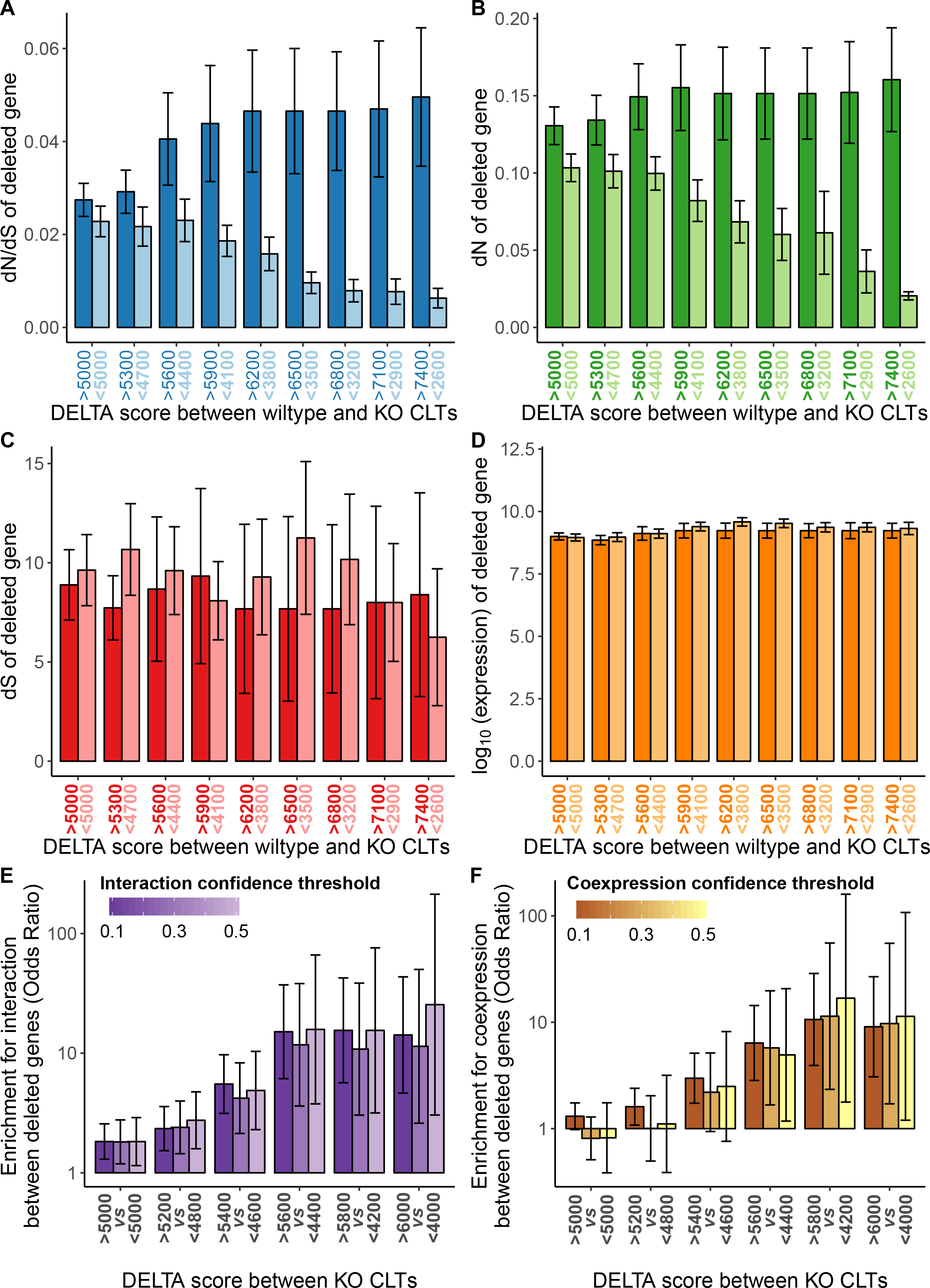
DELTA relates phenotypic changes in CLT to underlying genetic mechanisms (A-D) *C. elegans* single gene KO CLTs were categorized into high (dark color) or low (light color) DELTA score groups by different thresholds (*x* axis) according to their global alignments with wildtype CLT. For dN/dS (A) and dN (B), but not dS (B) or expression level (D) of the deleted genes, their differences becomes more dramatic as the DELTA score becomes more separated for the high and low group. Error bar indicates standard deviation assessed by 1,000 bootstraps of the genes. (E-F) Enrichment with experimentally determined interactions (E) and coexpression (F) in pairs of genes with KO CLTs of high DELTA score, relative to those with low DELTA score was assessed by odds ratio from Mantel-Haenszel Test, where the thresholds for high and low DELTA score are indicated by x axis. The experimentally determined interactions and coexpression relationship were extracted from the STRING database with different confidence thresholds, as indicated by the color scale. Error bar indicates 95% confidence interval of the odds ratio given by Mantel-Haenszel Test.

To better understand this observation, we compared dN and dS separately with the DELTA score. We found that difference in DELTA scores are predictive of deviation in dN (Fig. 6B), but not dS (Fig. 6C). Since dS is primarily determined by mutation rate, whereas dN is determined jointly by mutation rate and natural selection ^31^, these results suggest that DELTA score indeed captured phenotypic changes in CLT that is subject to natural selection acting on the function of the gene, instead of a mutational bias in favor of less important genes. In addition, gene expression level, a potential confounding factor that is correlated with both evolutionary conservation and functional importance of a gene ^30^, was found unrelated to DELTA score (Fig. 6D). Therefore, the correspondence between the evolutionary conservation of a gene and the impact of its deletion quantify by DELTA is not confounded by expression level of the gene..

We next asked whether comparison between CLTs of two KO strains can reveal functional relationship between the deleted genes, as has been shown by CLT comparison using methods other than DELTA ^32–34^. Since DELTA score quantifies phenotypic similarity between two CLTs, we hypothesized that if the DELTA score between two KO CLTs is higher, the genes deleted in these strains are more likely functionally related. To test our hypothesis, we assessed the enrichment for experimentally determined interactions as recorded by STRING ^35^ in pairs of genes with KO CLTs of high DELTA score, relative to those with low DELTA score. To avoid interdependence between gene pairs due to involvement of the same gene, we construct a 2×2 contingency table for each of the 204 genes separately by (i) whether its DELTA scores with other 203 KO CLTs are higher or lower than some thresholds (Fig. 6E, *x* axis), and (ii) whether the confidence of experimentally determined interactions between the pair of deleted genes in comparison surpasses certain confidence threshold (Fig. 6E, color scale). The 2×2 contingency tables for all genes were summarized by Mantel-Haenszel procedure to calculate a combined odds ratio to reflect the enrichment of interaction. Similar analysis was also done for coexpression between deleted genes (Fig. 6F). For both interaction and coexpression, we found that the odds ratio increases as the DELTA score difference between the two groups of gene pairs becomes larger, regardless the confidence threshold used. For example, the enrichment of experimentally determined interactions with confidence > 0.5 yielded an odds ratio of 25.5 when gene pairs with DETLA score > 6000 is compared to those with DELTA score < 4000. These results suggest that genes whose deletion yield similar phenotypic outcome for CLT tend to be functionally related. Altogether, our comparative analyses among KO and wildtype CLTs by DELTA successfully associate CLT phenotypes to the underlying genotypes.

### CLT alignment between species by DELTA hints at evolutionary correspondence between cell types

The diversity of cell types is a significant feature of multicellular organisms, yet how it evolves remains largely unexplored. Just as one needs to find orthologous genes between species, the study of cell type evolution is impossible without a mapping of cell types, or “cell type orthology” between different species. Traditionally, cell types from two species are considered “orthologous” according to structure- and/or function-based cell type definition, such as neuron or muscle cells. Recent technological development of single cell RNA-seq and other high-throughput approaches ^36–38^ have promoted research efforts in revisiting cell type definition by molecular similarities at transcriptome or epigenome level, such as in the Human Cell Atlas Project ^39^. However, inferring cell type orthology by structural/functional/molecular similarities may not be reliable because they could have emerged from phenotypic convergence ^15^, such as found for the striated muscles of vertebrates and *Drosophila melanogaster* ^40^. Alternatively, we hypothesized that the DELTA score between CLTs from two species with non-uniformly defined cell types should be maximized by a scoring matrix based on the actual correspondence between specific cell types.

To test our hypothesis, we compared the CLT of *Pellioditis marina* and *C. elegans*, whose cell type identities were previously defined by structure and function non-uniformly in the two species ^26^. By a greedy strategy (See Methods, Fig. 7A and Fig. S4), we optimized the scoring matrix such that the DELTA score from global alignments between the two CLTs is maximally higher than those from pairs of control CLTs created by relabeling all cells of a type as another random type. Intriguingly, the high matching scores in the optimized scoring matrix indeed hints at biologically reasonable correspondence between cell types from the two species (Fig. 7B). For example, the optimized matrix suggests that the cells labeled as “Muscle”, “Death”, “Intestinal” and “Germ” have exact matches between the two species. On the other hand, cells labeled as “Nervous System” in *P. marina* correspond to the cells labeled as “Neurons” and “Structural” (Neuronal structural) cells; and cells labelled as “Pharynx” in *P. marina* corresponds to “Gland”, “Epithelial”, “Muscle” and “Neuron” cells in *C. elegans*. Last but not least, the 30 cells in *P. marina* with “Other” fate are suggested by the optimized matrix as epithelial cells in *C. elegans*. Furthermore, the DELTA-g alignment based on the optimized scoring matrix revealed the detailed correspondence between terminal cells of the two species (Fig. 7C). Some evolutionary events of cell fate changes are clearly highlighted, such as an Epiderm cell in *P. marina* becomes apoptotic in *C. elegans* (Fig. 7D). Collectively, these results exemplified the comparative analysis of CLTs by DELTA, which offers novel insights into the evolution of cell types and CLTs.

**Fig. 7.**
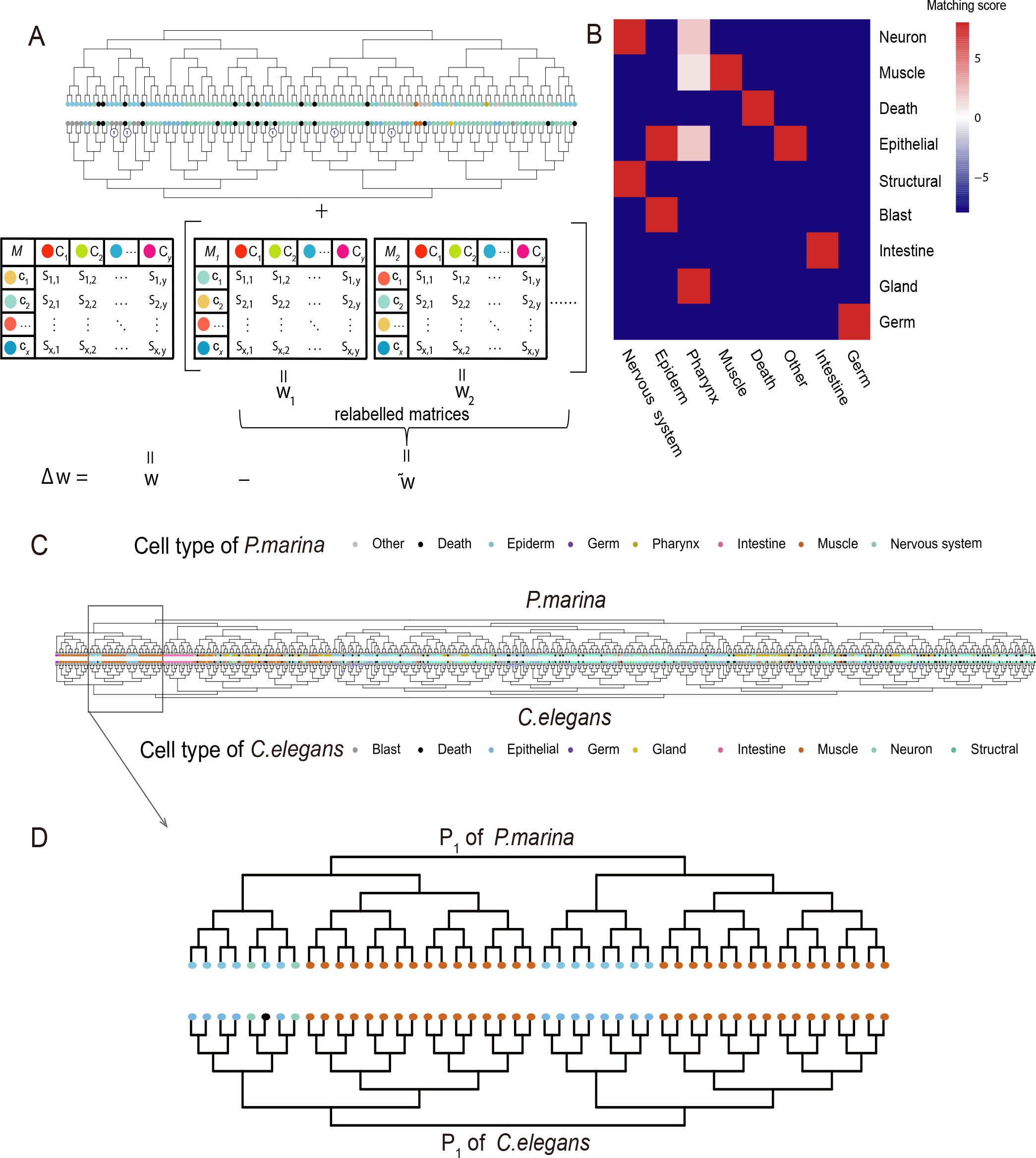
DELTA comparison across species highlights evolutionary correspondence of cell fates. (A) The optimization of scoring matrix between cell types of two species. With the cell types from two species, we first defined a scoring matrix *M*, whose row and column labels (cell types) were randomly permutated to generate control matrices *M*_1_, *M*_2_, … *M_i_*. These 1+*i* matrices were individually used by DELTA to align the CLTs from two species, giving rise to 1+*i* corresponding DELTA scores *w*, *w*_1_, *w*_2_, … *w*_*i*_. The deviation of *w* from its random expectation 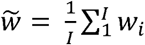, i.e. 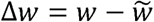, is optimized by a greedy strategy. See Methods and Fig. S4 for more details. (B) The optimal scoring matrix for comparison between *C. elegans* and *P. marina*. (C) The alignment between *C. elegans* and *P. marina* found by DELTA using the scoring matrix shown in (B). Note the small circled numbers on some internal branch indicate size of pruned sub-CLT. (D) A sub-alignment from (C), note the Epiderm “P1aabab” cell in *P. marina* is apoptotic in *C. elegans*.

## DISCUSSION

### A computational framework for comparative study of CLTs

In this study, we designed and implemented DELTA, a computational framework for the alignment of developmental cell lineage trees. Using simulated CLTs and real CLT from *C. elegans*, we showed that DELTA can find sub-CLTs with highly similar developmental programs, such as bilaterally symmetric lineage pairs, and lineages whose expression trajectory are highly correlated. Furthermore, DELTA alignments among mutant and wildtype *C. elegans* strains identify homeotic cell fate transformation, show more dramatic phenotypic change for the deletion of more important genes, and reveal higher CLT similarities between strains where functionally related genes were individually knocked out. Finally, we found that the scoring matrix, optimized for the DELTA score between CLTs from two species, can shed light on the evolution of cell types and CLTs of the two species. Together, our work has demonstrated the applicability of DELTA in multiple biological questions related to CLT, thus opening new paths to the analysis of CLTs, a type of data that will rapidly accumulate ^1,2,4,22,23^.

### Potential applications of DELTA

As more CLT being determined, the application of DELTA to them shall provide critical insight to several important biological questions. First, the repeatability of development can be assessed by comparing CLTs from different individuals, or CLTs that root at different single cells from an otherwise homogeneous cell population. This analysis is particularly relevant to the efficiency of iPS, where a seemingly homogeneous population of cells are similarly treated, but only a very small fraction are successfully transformed to the pluripotent state, with an even smaller fraction capable of growth into organoids. Comparison between the CLTs rooted at failed or successful iPS may lead to a mechanistic explanation for their differences.

Second, as demonstrated with simulated and wildtype *C. elegans* CLT, DELTA is capable of associating CLT phenotypes to the (epi-)genetic states of individual cells. Theoretically, the phenotypic consequence of (epi-)genetic states, or the genotype-phenotype mapping (GPM) shall become more complex when more cells were involved. As an intermediate phenotype between those of single cells and of tissues/organisms, DELTA offers a novel path of bridging GPM on unicellular and multicellular levels.

Third, we have shown in our study that DELTA can be used to find the evolutionary correspondence between cell types in two species. The advancement of single cell transcriptome profiling experiments has allowed cell type classification at its finest scale with molecular signatures of gene expression. Experimental pitfalls aside, this approach of cell type identity determination has two biological difficulties. On the one hand, similarity of transcriptional profile could have arisen from both cell type homology due to inheritance from common precursor, and phenotypic convergence, which might lead to false combination of different cell types into one. On the other hand, the stochastic nature of gene expression^41^ may lead to erroneous separation of homogeneous cell population of one type into two. This problem is also recently noticed ^15^, where evolutionary definition of cell types based on the “core regulatory complex” (CoRC) of transcription factors is proposed. As a complementary approach, DELTA utilizes the biological information in CLT to find the evolutionary correspondence between cell types, and simultaneously reveals how CLT itself evolves.

Fourth, one critical unanswered question in development is the relative prevalence of autonomous and regulative development. In nematodes such as *C. elegans*, development is autonomous except for a small number of sub-CLTs ^42^. In most other animals, however, it is generally believed that regulatory development is the prevailing mode and autonomy the exception. A direct quantitative answer to this question would emerge by examining the result from DELTA local alignments for the frequency of identical sub-CLTs, should CLT or sub-CLT be available for those species.

Fifth, an ideal CLT comprise complete longitudinal and horizontal data. However, during experimental assessment of CLTs by single cell transcriptome profiling and lineage barcoding, cell lyses and losses are inevitable in the state-of-the-art method, which respectively dictate that CLT is longitudinally (because the cells are killed at the time of experimentation) and horizontally incomplete. DELTA may just provide a resolution to this problem by allowing assembly of temporally “sliced” incomplete CLTs, just as sequence alignment had allowed assembly of the genome out of short reads.

### Limitations and future improvements

There are several potential caveats in our study that worth discussing. Apparently, DELTA result critically relies on the choice of its two parameters, the scoring matrix between cell types and the pruning cost. Although we’ve carefully chosen biologically informed parameters (See Methods), there has been no objective estimation on how good or bad they are, which is likely impossible before more CLT data becomes available, much like the refinement of substitution matrix for sequence alignment when more sequence was determined^43^. Also, the value of pruning cost relative to the matching score also affects DELTA results. On the one hand, lower pruning cost makes DELTA more sensitive, because pruning of small sub-CLTs improves alignment compared to terminal cell type mismatches. On the other hand, higher pruning cost makes DELTA more specific, since terminal cell type mismatches are more likely retained than pruned. Nevertheless, a poor choice of parameters likely reduces the biological signals. The significance of all patterns we’ve shown in the paper would thus be stronger if the parameters were further optimized, further enhancing the value of DELTA.

We have considered qualitative cell type or epigenetic states in our definition of CLT. However, the dynamic programming scheme for comparison between CLTs is readily adaptable to quantitative definition of cell types made by high throughput experiments such as single cell transcriptomics. In this case, instead of using the scoring matrix between qualitatively defined cell types, the alignment score between a pair of terminal cells will be calculated by the similarities between their transcriptome profiles, using quantitative metrics such as correlation coefficient or negated Euclidean distance.

In the current study, CLTs were used without considering the temporal duration of cell cycle for each cell. Apparently, discarding the information of cell cycle duration did not prevent us from discovering the similarities in developmental program by analyzing CLTs. There are two potential explanations for this observation. On the one hand, the molecular pathway that regulates cell cycles might be tightly coupled to the developmental program, such that cell cycle duration is an intrinsic property of the cell type. In other words, most cell types have their own specific cell cycle duration, so that the definition of cell types already contains the information of cell cycle duration. On the other hand, the definition of cell types might have nothing to do with the cell cycle duration, where DELTA needs to be further improved by allowing cell cycle duration adjustment as an additional CLT edition (besides sub-CLT pruning), with properly defined costs to DELTA score.

For both simulated CLTs and real CLTs from *C. elegans*, the developmental process is mostly autonomous, whereas in most complex organisms (except nematode), appears largely regulatory. The critical difference between autonomous and regulatory development is whether the epigenetic states of individual cells can be altered by external cues, such as environmental stress or signals from other cells.

For example, when isolated from the 8-cell stage mouse blastomere, one cell can grow into one individual mouse ^44^, but not one eighth of a mouse, as would be predicted by autonomous development. However, autonomous cell fate determination is certainly not absent in complex organisms, especially towards the end of the developmental process. Meiosis is one such example, where a primary oocyte divides twice (three division events) and creates one mature ovum and three polar bodies. As along as such autonomous sub-CLTs exist, DELTA alignment would be possible and informative, as demonstrated by our simulated CLTs with perturbed genes expression (Fig. 2).

Finally, current single cell high throughput experiments suffer from loss of cells, thus the CLT reconstructed by lineage tracing DNA barcode is likely incomplete, with majority of terminal cells missing. The robustness to such data quality issues dictates the applicability of DELTA. As shown in our DELTA analysis by simulated CLTs with randomly dropped terminal cells, although the alignment becomes smaller as more terminal cells are lost, those sub-CLT pairs aligned by DELTA remains highly similar in epigenetic state. In other words, the loss of terminal cells decreases sensitivity of DELTA, whereas the specificity likely remains high (Fig. 2C-G). If more CLT data were to become available, this limitation would also likely be alleviated, just as short sequence fragments with high enough coverage could be assembled into the full-length sequence.

Collectively, DELTA establishes a computational foundation for the alignment of CLTs and potentiates systematic analyses of lineage trees. Together with the accumulating experimental data of CLTs, DELTA will likely illuminate the connection between phenotypes represented by CLTs and their underlying genotypes, providing novel insights to many unresolved biological questions.

## Methods

### The DELTA algorithm

Given an alignment (not necessarily optimal) of two CLTs, two parameters are required to quantify how similar these two CLTs are. First, for any pair of cell types comprising one cell type from each of the two CLTs being compared, a “matching score” is required to describe their similarity in developmental state. All the matching scores between the cell types from one CLT and the cell types from the other CLT can be summarized as a scoring matrix, in analogy to the substitution matrix (e.g., PAM, BLOSUM) used in sequence alignment, except that cell types from the two CLTs could be different. Second, similar to gap penalty in sequence alignment, a pruning cost is used when some cells or sub-CLTs from one CLT is “pruned”, meaning no correspondence in the other CLT can be found for them. In our implementation, the pruning cost is multiplied by the size (number of leaves) of the pruned sub-CLTs, and then simply subtracted from the alignment score when sub-CLT pruning is required. With scoring matrix and pruning cost defined, the goodness of correspondence between terminal cell types and topology can be quantified as a score for the alignment between a pair of CLTs. The task of DELTA is then to find the alignment with the maximal possible score (the “DELTA score”), allowing necessary mismatches of types in terminal cells and pruning of sub-CLTs.

Finding the optimal alignment between CLTs is computationally intensive, because a CLT remains unchanged, or isomorphic, by swapping any pair of sister sub-CLTs. Thus a query CLT with 1,000 internal nodes, approximately the size of the full *C. elegans* CLT, could be aligned to another subject CLT in 2^1,000^ possible ways, not to mention the isomorphic transformation of the subject CLT, and pruning. In DELTA, this issue is resolved by the dynamic programming ^24,25,45^ (Fig. S1A), where the smallest sub-CLTs (those containing only one terminal cell) were aligned first, and larger sub-CLTs were aligned by the best combination of the alignments of its two daughter sub-CLTs, be it match, mismatch, or pruning. The final alignment is extracted from the dynamic programming matrix by backtracing the sub-alignments from the matrix cell with the top DELTA score.

To gauge the statistical significance of an alignment, for each of the aligned CLTs, we generated 1,000 pseudo-CLTs by randomly coalescing the leaves of the real CLT and calculated the DELTA score between the 1,000 pairs of pseudo-CLTs. (Details given in Supplementary Text S1). The distribution of DELTA score assessed by these randomized CLTs controls both the sizes of the CLT, as well as the composition of the terminal cells. The DELTA scores of the 1,000 pairs of pseudo-CLTs were used to estimate a *P* value for the actual DELTA score by *Z*-test. We implemented DELTA algorithm in C++, whose source is available on github. Three files are required as input for DELTA algorithm, including two files with trees to be aligned, and a file defining the matching scores of different terminal cell types (Fig. S3).

### Simulated CLT

We simulated CLTs using a previously published model ^27^. Briefly, the expression profile of a cell at time *t* is represented by vector *S*(*t*), whose elements *s*_*i*_(*t*) (−1 ≤ *s*_*i*_(*t*) ≤ 1) indicate the expression of genes *i* = 1,2,…,*N* (*N* > 2), and the epigenetic state of the gene is considered as “on” if *s*_*i*_(*t*) > 0, or “off” otherwise. A regulatory network composing of these *N* genes are constructed as a N × N matrix *R*, whose elements *r*_*ij*_ indicate the regulatory effect of gene *i* on the expression of gene *j* (Fig. 2A). We defined that each gene regulates an average of *K* other genes, and thus random values following standard normal distribution was assigned to *N* × *K* random elements in R, whose remaining elements were set to 0, indicating no regulation (Fig. S2A). Therefore, the expression profile of the cell at the next discrete time point *S*(*t*+1) can be calculate algebraically using *S*(*t*) and *R* as S(t+ 1) = f(R × S(t)), where 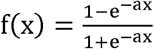 is a sigmoid activation function that determines how the expression of each gene is influenced by the total regulatory input from the interaction network ^46,47^, and *a* is the activation constant that determines the transition shape of the sigmoid curve. Among the *N* genes, two of them have special roles as cell cycle regulator and asymmetric division regulator, respectively. For the cell cycle regulator (gene 1), the cell divides instantaneously into two daughter cells once *s*_1_(*t*) > 0, and *s*_1_(*t*) for both daughter cells are reset to −1. For the asymmetric division regulator (gene 2), if the cell divides when *s*_2_(*t*) ≤ 0, both daughter cells retain the original expression of *s*_2_(*t*), otherwise, one of the daughter cells will be assigned *s*_2_(*t*) = −1, while the other daughter cell retains the original *s*_2_(*t*). The developmental process is then simulated by initializing a single cell with a randomly generated *S*(0), with which *S*(1), *S*(2), and so on, is calculated. Multiple rounds of cell division as dictated by the regulatory network will be recorded until *t* = *t*_max_ or the “depth” of any terminal cells reach *d*_max_, where “depth” refers to the number of cell divisions a terminal cell undergoes since the zygote. This procedure gives rise to a CLT, where the terminal cell types are defined by *S*(*t*_max_) (Fig. 2A). Note that *t*_max_ and *d*_max_ not necessarily indicate the end, but rather a “cross-section” of the full developmental process.

We performed 1,000 tree simulations for each condition, and performed self-alignment using a scoring matrix defined by the number of genes with identical epigenetic on/off states between a pair of cell types, and a pruning cost of 1.25*N*. In the main text, we showed results from *a* = 100, *N* = 16, K = 2, *t*_max_ = 50 and *d*_max_ = 6. We tried various different settings (Fig. S2B) and found our observation of DELTA’s capability of associating CLT with epigenetic similarity to be robust in different settings. For example, most of the aligned sub-CLTs have very small *P*-values (Fig. S2C), which means that this DELTA score is much higher than that between two random trees. Also, the log10(*P*-value) is highly correlated with the match length (Fig. S2C) and alignment scores (Fig. S2D) of the subtrees, indicating that the more complex two subtrees are, the lower chance that the alignments can be generated by chance.

CLT of other species might not be fully autonomous and deterministic as *C. elegans*, and experimentally determined CLT might not capture all terminal cells. We modeled these issues as two types of perturbations in CLT simulation. On the one hand, expression of each gene has a 5% probability to be negated at every time point in every cell. On the other hand, 35% of terminal cells were randomly removed, and an apparent CLT was reconstructed following the topology of the real underlying CLT. That is, any internal cells that lost all its descendant leaves were also removed, and those that loss one of its daughter cells were replaced by the remaining daughter.

### Experimentally determined cell lineage trees

The developmental cell lineage tree of wildtype *Caenorphabditis elegans* as determined by Sulston et al. ^16^, and that of *Pellioditis marina* as determined by Houthoofd et al. ^18^ were retrieved from previous publications ^7,26^. Briefly, the 671 terminal cells in *C. elegans* cell lineage (up to hermaphrodite embryogenesis) were categorized by standard anatomical descriptions ^16^ as: 39 blast, 113 death, 93 epithelial (arcade, hypodermis, pharyngeal structural, rectum, valves), 2 germ, 13 gland (coelomocytes, excretory system, and pharyngeal glands), 20 intestinal, 123 muscle (including the head mesodermal cell), 46 neural structural cells, and 222 neurons. DELTA comparison between *C.elegans* CLTs with anatomically defined terminal cell types was carried out using a scoring matrix where the cell pairs of identical types are scored by 10 and other pairs by −2, and a pruning cost of 1. For *P. marina*, the cell lineage with 638 terminal cells (up to muscle contraction) were classified as: 81 body muscle, 67 death, 2 germ, 131 hypodermis, 20 intestine, 195 nervous system, 112 pharynx, and 30 other fate ^18^. Bilaterally symmetric sub-CLTs in *C. elegans* were extracted from previous reports ^16^.

For gene expression along the *C. elegans* lineage tree, we downloaded the EPIC (Expression Patterns in Caenorhabditis) data ^29^. Of genes with more than one biological replicates, only the one used as examples on the website was used. For each individual gene, we further averaged its expression across the whole lifespan of each cell to generate its expression level for the individual cell. Each cell was then represented by the expression of all 130 genes recorded in EPIC, and one Pearson’s correlation coefficient was calculated by concatenating expressions from all aligned cells in one CLT, with that from the other CLT in the alignment. Since different CLT alignments involve different number of aligned cells, before the Pearson’s correlation coefficients (*R*) were standardized by 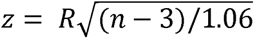 (Fisher’s r-to-z transformation) before being compared (Fig. 4).

To evaluate the capability of DELTA in associating phenotypic changes in CLT to their underlying genetic changes, we downloaded the CLT of *C. elegans* where 204 conserved genes were individually knocked down from the Digital Development database ^3^. The downloaded data contain the epigenetic state (“ON” or “OFF”) of three tissue markers, namely cnd-1 (a subset of neurons), pha-4 (pharynx and gut), nhr-25(HYP). Assuming cells with the same lineal name in different experiment are cells with the same identity, we further combined the experimental replicates for the same mutant CLT by a simple majority rule. That is, the epigenetic state of this marker in a specific cell is considered “ON” if it is supported by the majority of the experimental replications of the specific mutant strain. With the epigenetic states of the three markers, each terminal cell was categorized into one of eight (2^3^) types. To construct a scoring matrix as DELTA parameter, the matching score between two cell types were defined as 10*x* – *y*, where *x* and *y* are respectively the number of markers with same and opposite epigenetic states in the two cell types (Fig. 5A). The pruning parameter was set to 1. During DELTA comparison between two CLTs, it is possible that some terminal cells from one CLT become internal in the other CLT. To ensure their comparability, we removed any cell that is recorded in only one of the two CLTs based on their lineal names, and used the remaining ancestral internal cells (mother of the removed cells) as terminal. The list of homeotic transformation previously discovered was manually retrieved from the original report ^5^.

### Genomic and comparative genomic data

The expression level of *C. elegans* protein coding genes, and the number of synonymous (*dS*) and nonsynonymous (*dN*) substitutions between one-to-one orthologs in *C. elegans* and *C. briggsae* was obtained as previously described ^30^. The confidence score for experimentally determined protein-protein interaction and gene coexpression among *C. elegans* genes were extracted from the STRING database ^35^ v10.5.

### Optimizing scoring matrix between cell types from two CLTs

To find the proper scoring matrix for alignment between CLTs of two species, such that correspondence between subjectively defined cell types can be inferred from DELTA, we employed an Expectation Maximization algorithm to optimize the scoring matrix between cell types from two CLTs. The basic logic behind this algorithm is that a biologically meaningful scoring matrix should maximize 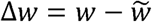, where *w* is the DELTA score from global alignment between the two CLTs being compared using a specific scoring matrix, and 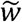 is the expected DELTA score when the same pair of CLTs is being compared using scoring matrix where cell types (labels of rows and columns) were randomly shuffled (Fig. 7A). For alignment between two CLTs with different cell types, say *x* and *y* respectively, there are *x* * *y* matching scores to be optimized. We employed a greedy grid search strategy to reduce the computational time of this optimization (Fig. S4). Briefly, the scoring matrix was initialized by assigning 0 to all elements, and then the scoring matrix was optimized by four rounds of grid search with increasing precision. Each round of grid search is finished by progressively optimizing multiple groups of 4 elements, with elements associated with more cells were optimized first (Fig. S4).

The details of the optimization are as follows. In the first round of grid search, for the first four element in the scoring matrix, we assigned −1 or 1 to each of the four elements, giving rise to 2^4^ different scoring matrices, and calculate 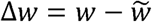 for each matrix, where 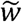 is the averaged DELTA score between the two CLTs by all the scoring matrices generated by permutating the cell types (column and row labels of the matrix) associated with the four elements being optimized (i.e., max number of permutation is 4! * 4! – 1= 575) (Fig. S4A). The one out of 2^4^ scoring matrices who has the largest ∆*w* was chosen. The next four elements were then optimized similarly. The whole scoring matrix was optimized by progressively optimizing groups of 4 elements until all elements were scanned once (Fig. S4B). In the second round of optimization, those elements assigned as −1 and 1 were further optimized for a choice between −2 or −1, and 1 or 2, respectively, by a method similar to the first round. The third round then continues, resulting in a scoring matrix whose elements were one of (−4,−3,−2,−1,1,2,3,4), where −4 and −3 were from elements valued −2 in the previous round, −2 and −1 were from −1, and so on. The final round of optimization will give rise to a scoring matrix whose elements were one of (−8,−7, −6,−5,−4,−3,−2,−1,1,2,3,4,5,6,7,8) (Fig. S4C). The pruning cost was fixed at 10 times the maximal possible matching score during this process. I.e. pruning cost is 10, 20, 40 and 80 for round 1, 2, 3 and 4, respectively. A high matching score between two cell types in this final scoring matrix indicates that the two cell types are closely related as suggested by the DELTA alignment between CLTs.

