## Supplementary Text 1 for "Computational comparison of developmental cell lineage trees by alignments"

**Supplementary Materials for “Computational comparison of developmental cell lineage trees by alignments”**

**Supplementary Text 1**

**Supplementary Figure S1**

**Supplementary Figure S2**

**Supplementary Figure S3**

**Supplementary Figure S4**

**Supplementary Text 1**

**Algorithmic details of DELTA**

A cell lineage tree (CLT), as used in the current study, can be described as a directed, acyclic, connected graph $T=\left( V, E \right)$, where *V* and *E* are collection of all nodes (vectors) and edges in the tree, respectively. Each node $v\in V$ represents one single cell, and each edge $e\in E$ represents a descendant relationship pointing from a mother cell to one of its daughter cells. The number of edges attached to a node (regardless the direction) is called the degree of the node, denoted by *d_v_*. The root node of the CLT has *d_v_* = 2, indicating the common ancestor of all the cells in the CLT, e.g. the zygote that gives rise to the whole *C. elegans* CLT. An internal node of the CLT has *d_v_* = 3, indicating non-root cells that undergoes further division recorded by the CLT. Note that we consider CLT as an unordered tree, which means that swapping the two daughters of an internal node, along with their descendant subtrees, does not change the tree. A terminal node of the CLT has *d_v_* = 1, which represents a terminal cell recorded by the CLT, but not necessarily the terminal of development (i.e., further divisions of the “terminal” cells are just not recorded as part of the CLT). All terminal nodes are labeled by their cell types, which could be anatomically defined as, for example, muscle, neuron, or epigenetically defined such as CD4+ cells.

DELTA finds alignments between two trees with necessary pruning of some subtrees. For a tree $T=\left( V, E \right)$, a subtree $T_{v}^{u}$ is a connected subgraph of *T*. $T_{v}^{u}$ contains all nodes $v'\in V$ such that the path between *u* and *v’* starts with the edge $\left( u\to v \right)$, as well as all the edges attached to these nodes except $\left( u\to v \right)$. A pruning of subtree $T_{v}^{u}$ includes three steps. First, remove all the nodes and edges in $T_{v}^{u}$; second, if *u* has two remaining neighbors attached to it by $\left( u'\to u \right)$ and $\left( u\to u'' \right)$, connect them by a new edge $\left( u'\to u'' \right)$; third, remove *u* and all edges attached to it.

Given a pair of trees $Q=\left( V,E \right)$ and $S=\left( V',E' \right)$, an isomorphic alignment is a bijection *A* : $V ⟷V'$, such that for every pair of nodes with $v,u\in V$, we have $\left( v,u \right)\in E\Leftrightarrow\left( A\left( v \right),A\left( u \right) \right)\in E’$. A homeomorphic subtree alignment *A* between *Q* and *S* is defined as an isomorphic alignment between $Q'$ and $S'$’, where $Q'$ is the result of zero or more pruning of subtrees in *Q*, and $S'$ is the result of zero or more pruning of subtrees in *S*. Here all the subtree pruning in *Q* and *S* are collectively denoted as $\pi\left( A \right)$. If we further denote the alignment score between two nodes $v\in V$ and $v'\in V'$ as $a\left( v,v' \right)$, and the cost for pruning a subtree $\hat{T}$ as $p\left( \hat{T} \right)$. The score of a homeomorphic subtree alignment *A* between Q and *S* can then be expressed as

$$w\left( Q,S,A \right)=\sum_{\left( v,v' \right)\in A} a\left( v,v' \right)-\sum_{\hat{T}\in\pi\left( A \right)} p\left( \hat{T} \right)$$

Given two CLTs, a scoring matrix *M* and a pruning coefficient *q* (>0), the DELTA algorithm finds the optimal *A* (with optimal/highest possible *w*) by dynamic programming (detailed below). Here, $p\left( \hat{T} \right)$ equals to *q* times the number of terminal nodes in $\hat{T}$. If $v$ and $v'$ are internal nodes, $a\left( v,v' \right)$ equals 0, or if $v$ and $v'$ are terminal nodes, $a\left( v,v' \right)$ equals to $M(f_{v},f_{v'})$, with $f_{v}$ and $f_{v'}$ representing the cell type of $v$ and $v'$. To find the optimal *w*, DELTA employs a modified implementation of previously described HSA algorithm (Milo et al., 2013) that was used to aligned RNA structure trees, with simplifications tailored for the alignment of CLT.

The dynamic programming (DP) procedure by which DELTA finds the optimal alignment between two CLT (*Q* and *S*) by recursively finding the optimal alignment between their subtrees. It starts from constructing a DP matrix with $N_{Q}$ row and $N_{S}$ column, where $N_{Q}$ and $N_{S}$ are the total number of nodes in *Q* and *S*, respectively. Each cell of the matrix will store the optimal *w* between the nodes/subtrees represented by the row and the column. For simplicity, we will hereinafter refer to optimal *w* between CLT *Q* and *S* as *w*(*Q*,*S*), with the optimal homeomorphic subtree alignment *A* implicitly indicated. To fill up the matrix, the *w* between terminal nodes are first directly determined by the scoring matrix of the terminal cell types. To calculate *w* between (i.e. to align) one leaf *v* and one internal node *u* with two daughter cells *l* and *r* (which could be internal nodes or leaves), one of the two subtrees ($T_{l}^{u}$ or $T_{r}^{u}$) need to be pruned. Thus, we have $w\left( v,u \right)= max\left( w\left( v,l \right)-p\left( T_{r}^{u} \right), w\left( v,r \right)-p\left( T_{l}^{u} \right) \right)$, where the optimal choice (indicating how should *v* and *u* be aligned) was stored for later traceback (Fig. 1D, top panel). For the DELTA score between two internal nodes *v* and *u*, whose children are respectively *l*/*r* and *l’*/*r’*, we have

$$w\left( v,u \right)=max\left\{ \begin{aligned} w\left( l,l' \right)+w\left( r,r' \right) \\ w\left( l,r' \right)+w\left( r,l' \right) \\ w\left( v,l' \right)-p\left( T_{r'}^{u} \right) \\ w\left( v,r' \right)-p\left( T_{l'}^{u} \right) \\ w\left( l,u \right)-p\left( T_{r}^{v} \right) \\ w\left( r,u \right)-p\left( T_{l}^{v} \right) \end{aligned} \right.$$

Again, the optimal choice was stored for later traceback (Fig. 1D, middle panel). This process went recursively until the whole DP matrix were filled. For the global alignment between *Q* and *S*, the traceback procedure starts from *w*(*Q*,*S*), i.e., the one DP matrix cell representing the alignment between the roots of *Q* and that of *S*. The optimal alignment for each node was then determined by the optimal choice recorded in each DP matrix cell along the (branched) route of recursive traceback. For the local alignment, the one DP matrix cell with the highest *w* (not necessarily involving the root of CLT) was located, and the local alignment was extracted by recursive traceback starting from this very DP matrix cell. All the DP matrix cells on the route of back tracing were marked as “used”, and the highest *w* in the unused DP matrix cells will similarly be used to extract the second local alignment result. More local alignments can be found by repetitively locating the highest *w* in the progressively smaller sets of unused DP matrix cells.

In assessing the statistical significance of a global or local alignment, the two aligned CLT (or subtrees, in case of local alignment) were individually randomized and realigned by DELTA for 1,000 times. The 1,000 resulting DELTA scores were used to estimate a *P* value for the actual DELTA score by Z-test. The randomization of an individual CLT is conducted as follows. First, all the terminal cells of the original CLT were extracted from the tree to form a list of cells. Second, two cells were randomly chosen from the list and paired up as sister cells, creating a subtree represented by their (arbitrarily constructed) mother cell. Third, these two chosen cells were then removed from the list and replaced by their mother cell. Fourth, the second and their steps were repeated until only one cell is left in the list, thus creating a randomized CLT with the same terminal cells as the original CLT. The distribution of DELTA score assessed by these randomized CLTs thus controls both the sizes of the CLT, as well as the composition of the terminal cells.

### Supplementary figure S1

Fig. S1 More on DELTA algorithm

(**A**) Detailed example of DELTA alignment in Fig. 1B. Two CLTs, *Q* and *S*, with types of their terminal cells were color-labeled. Note the nontriviality of the problem when the correspondence between cells in *Q* and *S* is sought after, regardless the extreme simplicity of both trees. The two CLTs, plus a scoring matrix describing how the correspondence between specific cell types should be awarded, based on hypothetical biological similarities of each pair, and a pruning cost, are used as input. The DELTA algorithm then uses a dynamics programming (DP) strategy to search for the optimal alignment between the two CLTs. Some critical steps of the DP procedure is exemplified here. First, a DP matrix was constructed and filled by comparing smallest sub-CLTs (those containing only one terminal cell) first, as shown by the light blue-shaded elements of the DP matrix. Each score between an internal node and a terminal node (green shaded elements) was then calculated by choosing the best combination of pruning and matching, with the source of best scored saved for each comparison (top panel). Similar procedure was carried out for scores between two internal nodes (pink shaded elements), allowing (mis-)matches for both daughter cells or necessary pruning (middle panel). In both top and middle panels, all possible combinations of daughter cell alignments and the corresponding score were listed as arrows and scores with different colors, whereas the best score is always colored blue. Arrows with dashed lines indicate pruning is involved, or solid lines otherwise. The high-scored element in the DP matrix was used as a starting point for backtracing, where the optimal alignment between *Q* and *S* were represented by the backtraced route (gray arrows in bottom panel). The final result gives the optimal alignment between *Q* and *S*. Note that the aligned terminal cells were vertically matched, and the aligned internal cells were indicated by double-head arrows. (**B**) The actual DP matrix generated by a DELTA alignment of the *Caenorhabditis elegans* CLT of standard anatomical terminal cell type annotation, with an isomorphic version of itself, where 30% randomly chosen sister sub-CLT pairs were swapped. Each element of the DP matrix was colored from low (blue) to high (red) by the scored percentile, as indicated by the color scale bar on top. The bottom right corner was zoom-in for finer details as an inset. The bar-plot on top shows the number of leaves for the sub-CLTs represented by the column of the DP matrix. (**C**) The top local alignments found by DELTA comparing *C. elegans* CLT with itself. The terminal cell types were color-labeled as indicated by the legend at the bottom right corner. Note that except the first alignment of full CLT root *vs* root, each pair of aligned sub-CLTs appeared twice as both X *vs* Y and Y *vs* X, which is an expected behavior of local alignment of a CLT *vs* itself.

### Supplementary figure S2

Fig. S2 More on DELTA analyses with simulated CLT.

(**A**) A matrix representing the regulatory network in Fig 2A, which is used to simulate CLTs presented in Fig. 2B. (**B**) Other parameters we tried to simulate CLTs. Each row represent a parameter combination (See Methods), the last three columns indicate the fraction of self-alignments of the simulated CLTs with a *P* value below three different thresholds. (C and D) 1,000 CLTs simulated with the parameter set (a = 100, N = 16, K = 2, *t*_max_ = 50 and *d*_max_ = 12) were self-aligned by DELTA-l. The top 100 CLT alignments were extracted from each DELTA-l run. The relationship between the statistical significance of the alignment and the number of matched cell (C) or DELTA score (D) is shown. Linear models fitted with the 100,000 points are represented by the red lines in both (C) and (D).

### Supplementary figure S3

Fig. S3. Implementation of DELTA. (A) Input and output files for DELTA. Input files required for DELTA include three tab-delimited text files, including two for the CLTs to be aligned, and one for the scoring matrix. A truncated example output file is also shown. The lineal positive of each cell were represented as binary strings. (B) Usage of the DELTA program, as shown when run without any argument. (C) The command line of a demo run of DELTA, with the first few lines of the result files displayed.

### Supplementary figure S4

Fig. S4. Greedy strategy used in optimization of the scoring matrix for *C. elegans* vs *P. marina*. (A) Cell types containing more cells were arranged to the top left corner of the scoring matrix. Then elements in the matrix were optimized as sub-groups of *n* elements (we used *n* = 4 in this study). Only row and column labels related to the elements currently being optimized were permutated to generated control relabeled matrices, considerably reducing the computational time for $\tilde{w}$. (B) Elements were optimized in groups as in (A) starting from the top-left (associated with more cells) to the bottom-right (associated with less cells), prioritizing those on the top-left to bottom-right diagonal line. (C) The scoring matrix were optimized following the scheme of (B) in four rounds, where each element of the matrix was chosen from two diverging options conditioned on the result of the previous round, such that $\Delta w$ is maximized.
