## Supplementary figures and images for "Computational comparison of developmental cell lineage trees by alignments"

### Supplementary Figure S1

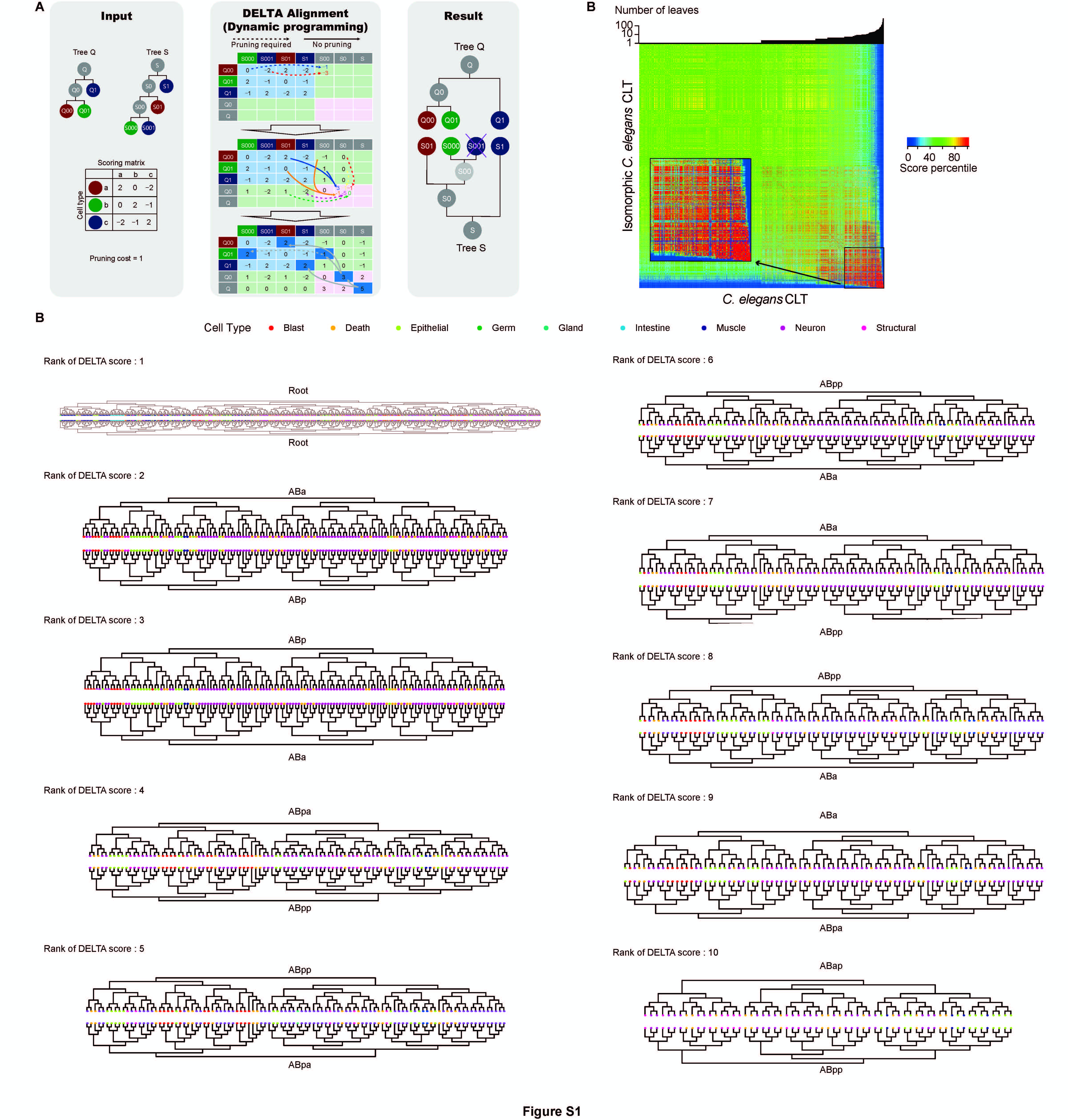

### Supplementary Figure S2

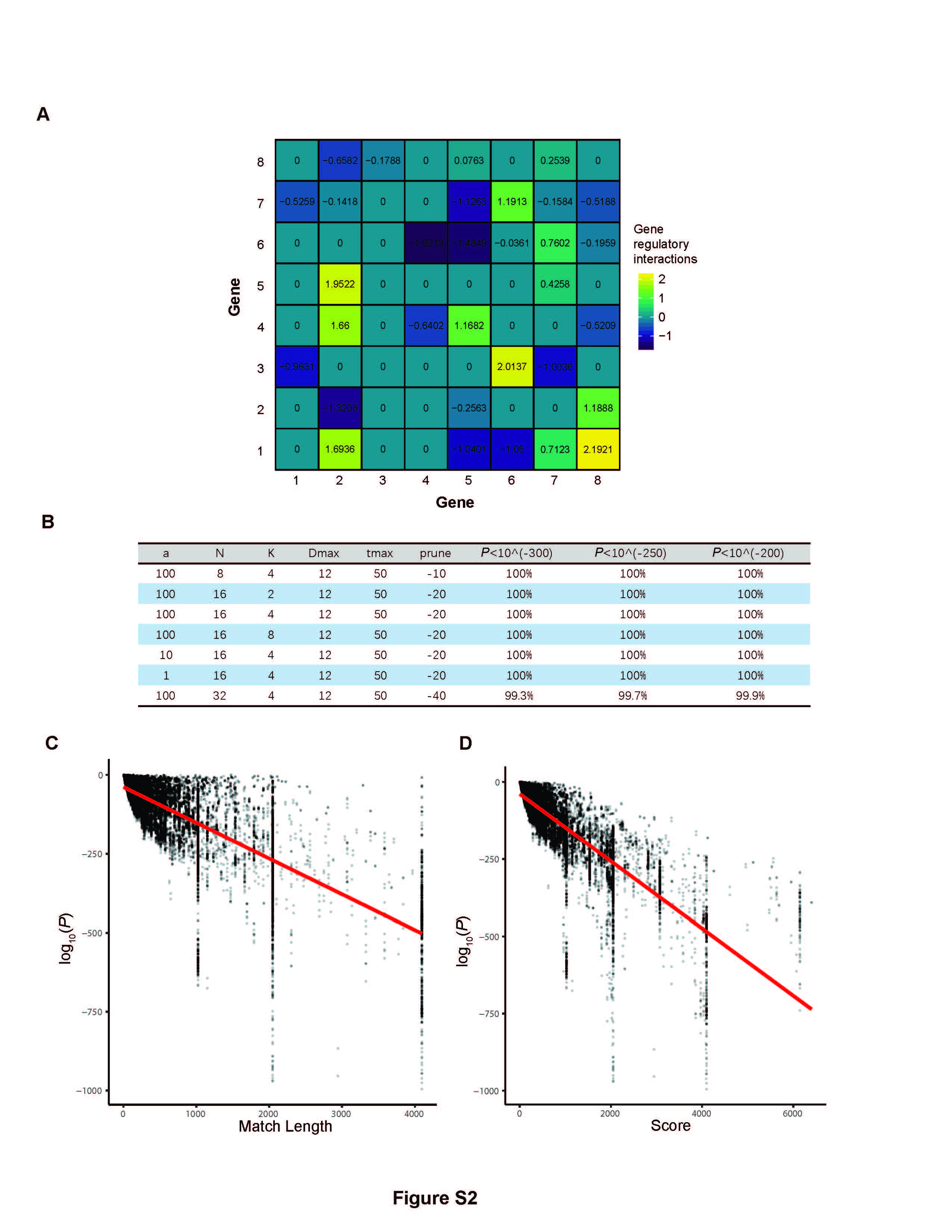

### Supplementary Figure S3

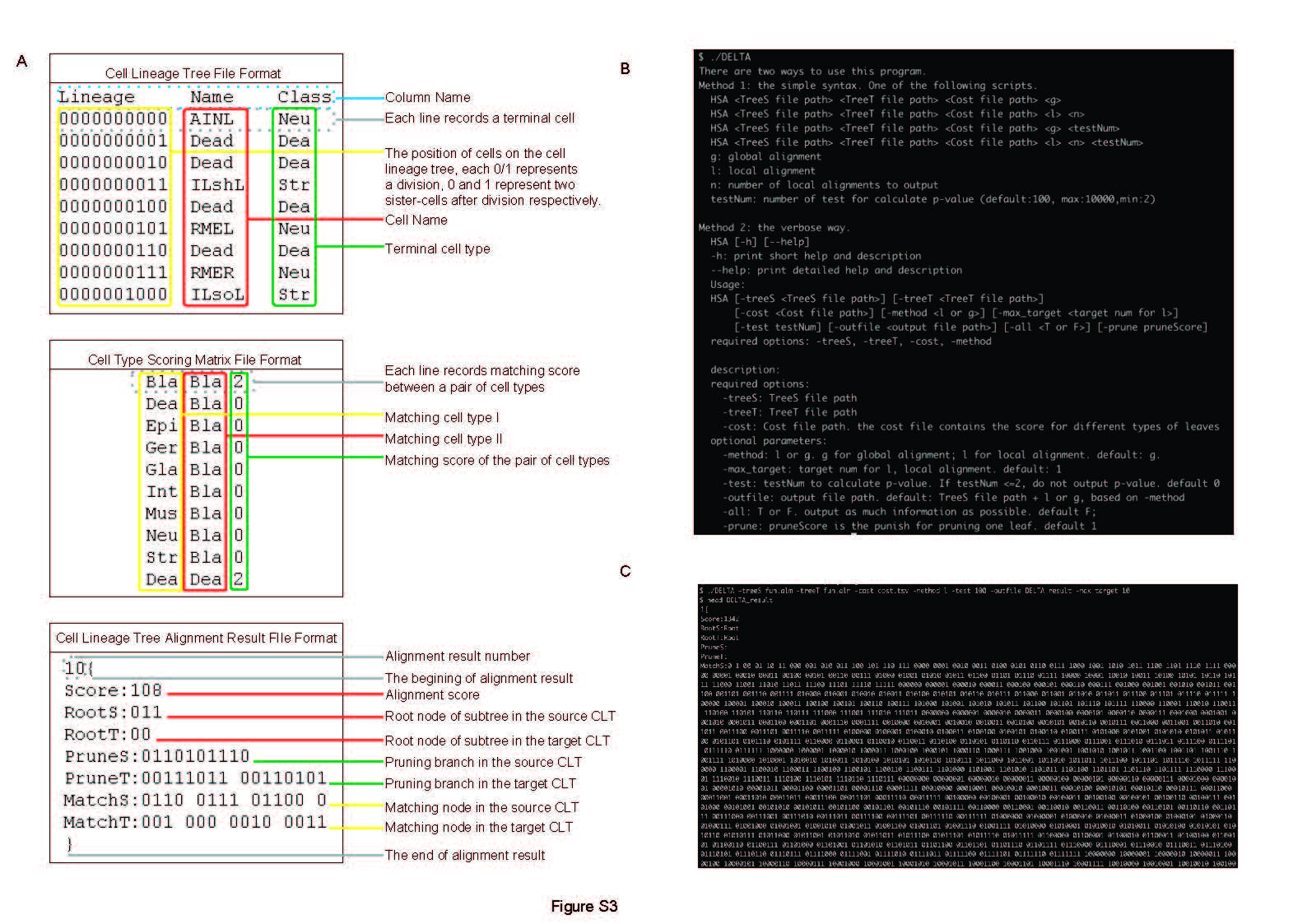

### Supplementary Figure S4

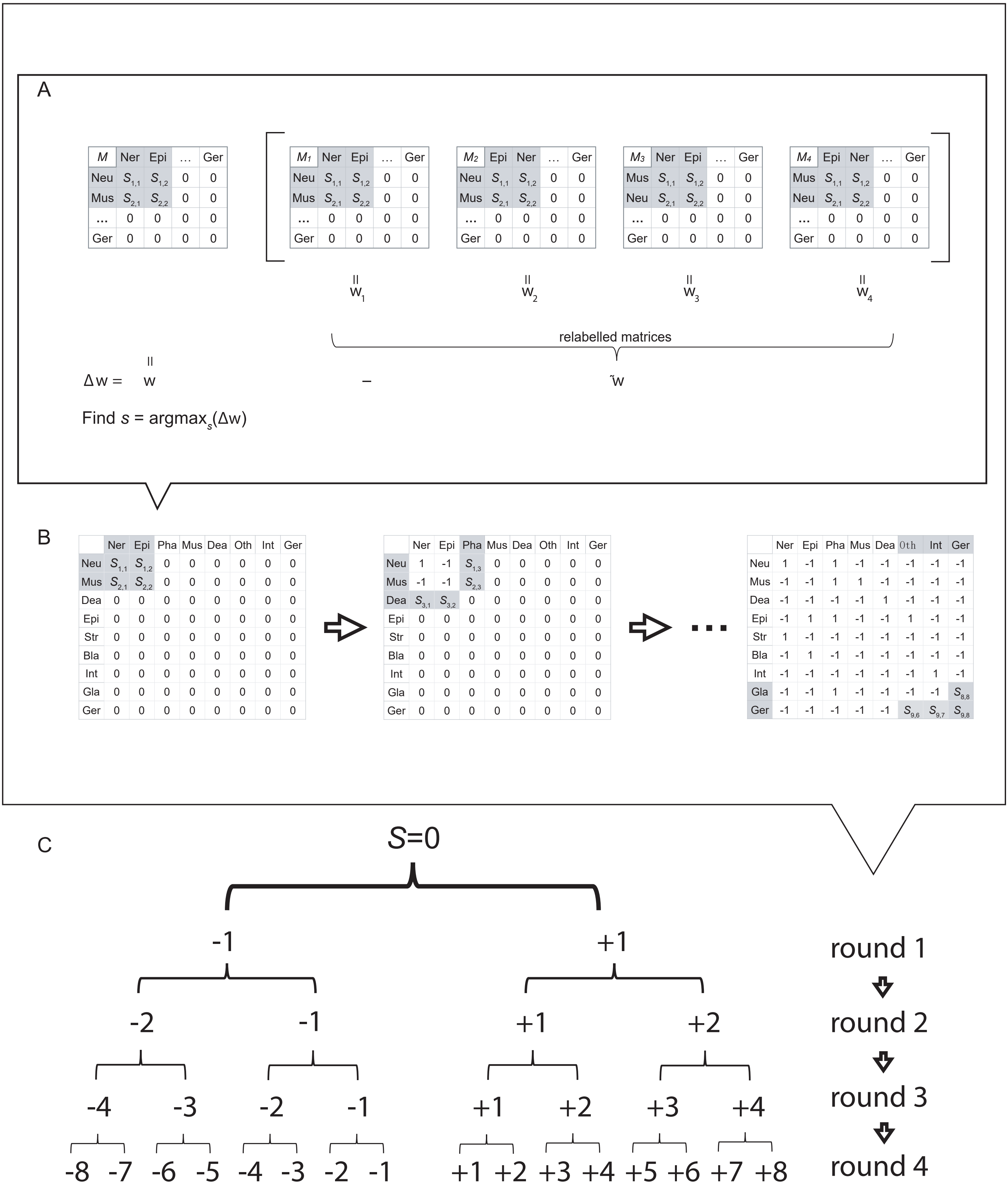
